# Concomitant DNA hydroxymethylation and histone H2B O-GlcNAcylation are prerequisites for zygotic genome activation in mice

**DOI:** 10.64898/2026.04.23.720328

**Authors:** Toshinobu Nakamura, Asuka Furuta, Tsunetoshi Nakatani, Toru Nakano

**Affiliations:** Nagahama Institute of Bio-Science and Technology, Shiga, 526-0829, Japan; JST, CREST, Saitama, 332-0012, Japan; Institute of Epigenetics and Stem Cells (IES), Helmholtz Zentrum München D-81377 München, Germany; Department of Pathology, Graduate School of Medicine, Osaka University, Osaka, 565-0871, Japan; Graduate School of Frontier Biosciences, Osaka University, Osaka, 565-0871, Japan

## Abstract

The paternal chromatin undergoes extensive reprogramming, characterized by the loss of 5-methylcytosine (5mC) through Ten-eleven translocation 3 (Tet3)-mediated oxidation to 5-hydroxymethylcytosine (5hmC). Given the role of DNA methylation in gene silencing, it has long been posited that this loss of paternal DNA methylation facilitates zygotic genome activation (ZGA), which occurs predominantly on the paternal genome. However, recent evidence indicates that Tet3-mediated 5mC oxidation alone does not influence global transcription in zygotes, leaving the molecular mechanisms governing ZGA largely elusive. Here, we identify the functional significance of O-linked N-acetylglucosamine (O-GlcNAc) modification of histone H2B at Serine 112 (H2BS112GlcNAc), catalyzed by O-GlcNAc transferase (OGT), within the paternal chromatin of mouse zygotes. We demonstrate that OGT is selectively recruited to the paternal chromatin because Stella (also known as PGC7 or Dppa3) inhibits its binding to the maternal chromatin—a recruitment mechanism analogous to that of Tet3. Although Tet3 and OGT associate with the paternal chromatin independently, both Tet3-dependent 5hmC formation and OGT-mediated H2BS112GlcNAcylation are indispensable for successful ZGA. Together, our findings reveal that a dual epigenetic signature—the simultaneous coordination of DNA hydroxymethylation and histone O-GlcNAcylation by Tet3 and OGT—is essential for initiating transcriptional reprogramming during the maternal-to-zygotic transition.

## Introduction

Zygotic genome activation (ZGA) is a critical nuclear reprogramming event that transitions the genome from a quiescent state to robust transcriptional activity shortly after fertilization. In mice, transcription initiates during the mid-1-cell stage and significantly increases at the 2-cell stage, periods referred to as minor and major ZGA, respectively (Li et al. 2013; Jukam et al. 2017; Svoboda 2018). This stepwise activation is essential for establishing totipotency and ensuring subsequent embryonic development. Despite extensive evidence highlighting the importance of ZGA, the underlying molecular mechanisms remain poorly understood. Given the dynamic chromatin remodelling during this period, epigenetic regulation likely plays a pivotal role in ZGA.

Previous studies have shown that Ten-Eleven Translocation 3 (Tet3)-mediated conversion of 5-methylcytosine (5mC) to 5-hydroxymethylcytosine (5hmC) induces a global loss of paternal 5mC (Iqbal et al. 2011; Wossidlo et al. 2011). Concurrently, the transcriptional activity of the paternal genome is significantly higher than that of the maternal genome (Bouniol et al. 1995; Aoki et al. 1997). Since DNA methylation typically functions as a transcriptional repressor, it was hypothesized that the loss of paternal 5mC is linked to minor ZGA. However, Tet3-knockdown zygotes—in which paternal DNA methylation is maintained—still initiate minor ZGA, indicating that the loss of 5mC itself is not required for this process (Inoue et al. 2012; Shen et al. 2014). Meanwhile, recent studies have revealed that Tet2/3 can function as scaffolding proteins that recruit O-linked N-acetylglucosamine (GlcNAc) transferase (OGT) to chromatin, leading to the GlcNAcylation of histone H2B at Serine 112 (H2B S112GlcNAc) (Chen et al. 2013). This Tet-OGT interaction is thought to promote transcriptional activation, presumably by influencing histone H3 lysine 4 trimethylation (H3K4me3) and H2B lysine 120 monoubiquitination (H2BK120ub) (Chen et al. 2013; Deplus et al. 2013). Since OGT-mediated H2B S112GlcNAc facilitates H2BK120ub and promotes a transcriptionally permissive chromatin state in embryonic stem (ES) cells (Vella et al. 2013), we hypothesized that this modification similarly contributes to ZGA. In this study, we de monstrate that although Tet3 and OGT bind independently to the male pronucleus, both Tet3-mediated 5hmC generation and OGT-mediated H2B S112GlcNAcylation are essential for successful ZGA.

## Results and Discussion

### Paternal-specific H2B S112 O-GlcNAcylation is regulated by Stella in zygotes

Since Tet3 is highly expressed in early stage of preimplantation embryos, we first examined whether chromatin contains H2B S112GlcNAc in the pronuclei stage zygote. As shown Fig. 1A, H2B S112GlcNAc was detected in paternal but not in maternal chromatin of pronuclei stage zygote. In contrast, both the maternal and paternal pronuclei were stained positively with anti-OGT antibody (Fig. 1B).

**Figure 1.**
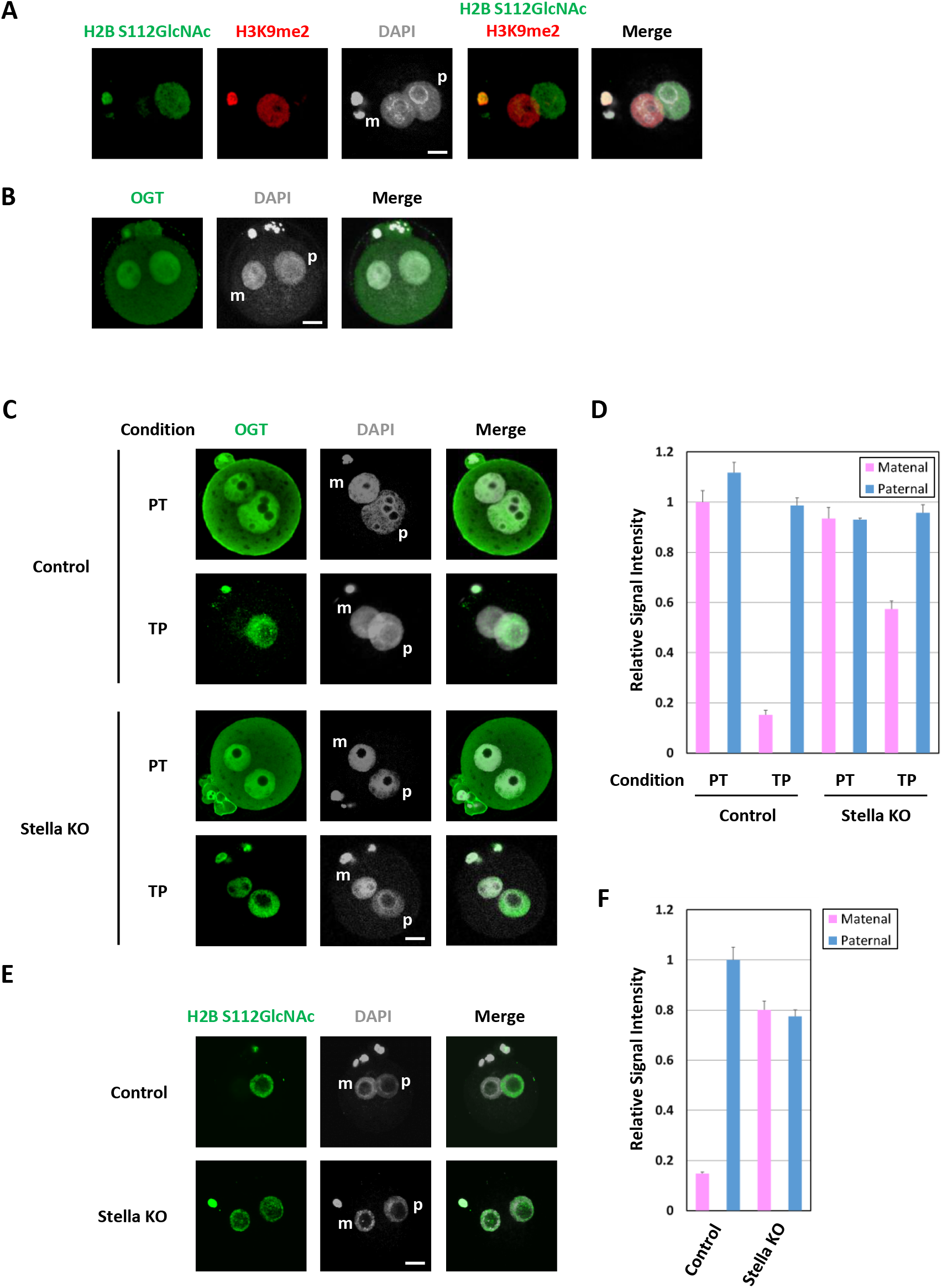
Paternal chromatin specifically contains H2B S112GlcNAc in zygote.t. (A) Zygotes were stained with the indicated antibodies (m, maternal pronuclei; p, paternal pronuclei). H2B S112GlcNAc and H3K9me2 are shown in green and red, respectively; nuclei were stained with DAPI (grey). (B) Zygote were stained with the anti-OGT antibody (m, maternal pronuclei; p, paternal pronuclei). OGT is shown in green; nuclei were stained with DAPI (grey). (C, D) Zygote obtained from female control or Stella KO mice were treated with PT or TP conditions as shown in Supplementary Fig. 1. Localization of OGT was analysed using anti-OGT antibody (m, maternal pronuclei; p, paternal pronuclei). OGT is shown in green; nuclei were stained with DAPI (grey) (C). Twenty-nine and 43 control zygotes were stained under PT and TP conditions, respectively. Nine and 16 zygotes obtained from Stella KO were stained under PT and TP conditions, respectively. Intensities of nuclear OGT signals of parental chromatin were quantified using Image J (D). Error bars indicate standard deviation. (E, F) Zygote obtained from female control or Stella KO mice were stained using anti-H2B S112GlcNAc antibody (m, maternal pronuclei; p, paternal pronuclei). H2B S112GlcNAc is shown in green; nuclei were stained with DAPI (grey) (E). Thirty-eight control zygotes and 11 zygotes obtained from Stella KO female mice were stained using anti-H2B S112GlcNAc antibody. Intensities of H2B S112GlcNAc signals of parental chromatin were quantified using Image J . Error bars indicate standard deviation. Scale bar, 20 μm.

We next examined the chromatin-binding characteristics of OGT by treating zygotes with Triton X-100 prior to fixation. As we previously reported (Supplementary Fig. 1) (Nakamura et al. 2012), this method allows free or loosely associated proteins to be eluted from chromatin. OGT was detected in both maternal and paternal chromatin after conventional paraformaldehyde (PFA) fixation (the PFA–Triton (PT) condition in Fig. 1B-D). In contrast, the staining pattern was markedly different when Triton X-100 treatment was performed before PFA fixation (the Triton–PFA (TP) condition in Fig. 1B-D). Under the TP condition, OGT was detected only in the paternal pronucleus (Fig. 1C, D). These results indicate that OGT is tightly bound to paternal chromatin.

We previously reported that the maternal factor Stella (also known as PGC7 or Dppa3) inhibits Tet3 function by binding to maternal chromatin enriched in dimethylated histone H3 lysine 9 (H3K9me2) (Nakamura et al. 2012). In addition, chromatin-bound Stella suppresses DNA methyltransferase 1 (Dnmt1) through Np95 (also known as Uhrf1 or ICBP90) in Stella-overexpressing somatic cells (Funaki et al. 2014), and it also inhibits Dnmt3a during iPS cell generation (Xu et al. 2015). Thus, although the specific targets vary depending on cellular context, chromatin-bound Stella generally acts as an inhibitor of chromatin modifiers. These findings prompted us to investigate whether Stella affects OGT binding to chromatin in zygotes. Under the TP fixation condition, OGT was detected not only in the paternal pronucleus but also in the maternal pronucleus of Stella-KO zygotes, demonstrating that Stella inhibits OGT binding to maternal chromatin (Fig. 1C, D). Consistent with this observation, H2B S112GlcNAc was detected in both maternal and paternal chromatin in Stella-KO zygotes (Fig. 1E, F). These data clearly indicate that H2B S112GlcNAc is suppressed by Stella in the maternal pronucleus.

### Tet3-dependent DNA hydroxymethylation occurs independently of OGT-mediated H2B S112 O-GlcNAcylation

Tet1 and Tet2 have been identified as OGT-binding partners and form independent high-molecular-weight nuclear complexes with OGT in mouse embryonic stem (ES) cells (Vella et al. 2013). Reciprocally, OGT was reported as an abundant interacting protein in Tet2- and Tet3-overexpressing HEK293T cells (Chen et al. 2013). We also confirmed the interaction between Tet3 and OGT in HEK293T cells by co-immunoprecipitation (Supplementary Fig. 2). Because more than 1,000 genes are known OGT targets, OGT knockout or knockdown approaches were not suitable for analyzing the function of H2B S112GlcNAc. Therefore, we employed an H2B S112A mutant, in which serine 112 is replaced with alanine, to examine the function of H2B S112GlcNAc in zygotes.

To elucidate the relationship between Tet3 and OGT in zygotes, we first attempted to deplete H2B S112GlcNAc by injecting mRFP-tagged H2B or its S112A mutant cRNAs into MII oocytes before fertilization, and subsequently obtained zygotes by *in vitro* fertilization (Fig. 2A). Both mRFP-H2B and mRFP-H2B S112A were efficiently incorporated into maternal and paternal chromatin (Fig. 2B, C; Supplementary Fig. 3). Expression of wild-type H2B did not alter H2B S112GlcNAc levels, whereas expression of the H2B S112A mutant resulted in complete loss of H2B S112GlcNAc in zygotes. Notably, 5hmC levels were unchanged in zygotes depleted of H2B S112GlcNAc, indicating that Tet3-mediated conversion of 5mC to 5hmC is independent of OGT-mediated H2B S112GlcNAc (Fig. 2B, C). Because Tet3 is reportedly recruited to chromatin by GSE (gonad-specific expression) in zygotes (Hatanaka et al. 2017), we cannot exclude the possibility that OGT is also recruited to chromatin by GSE in zygotes.

**Figure 2.**
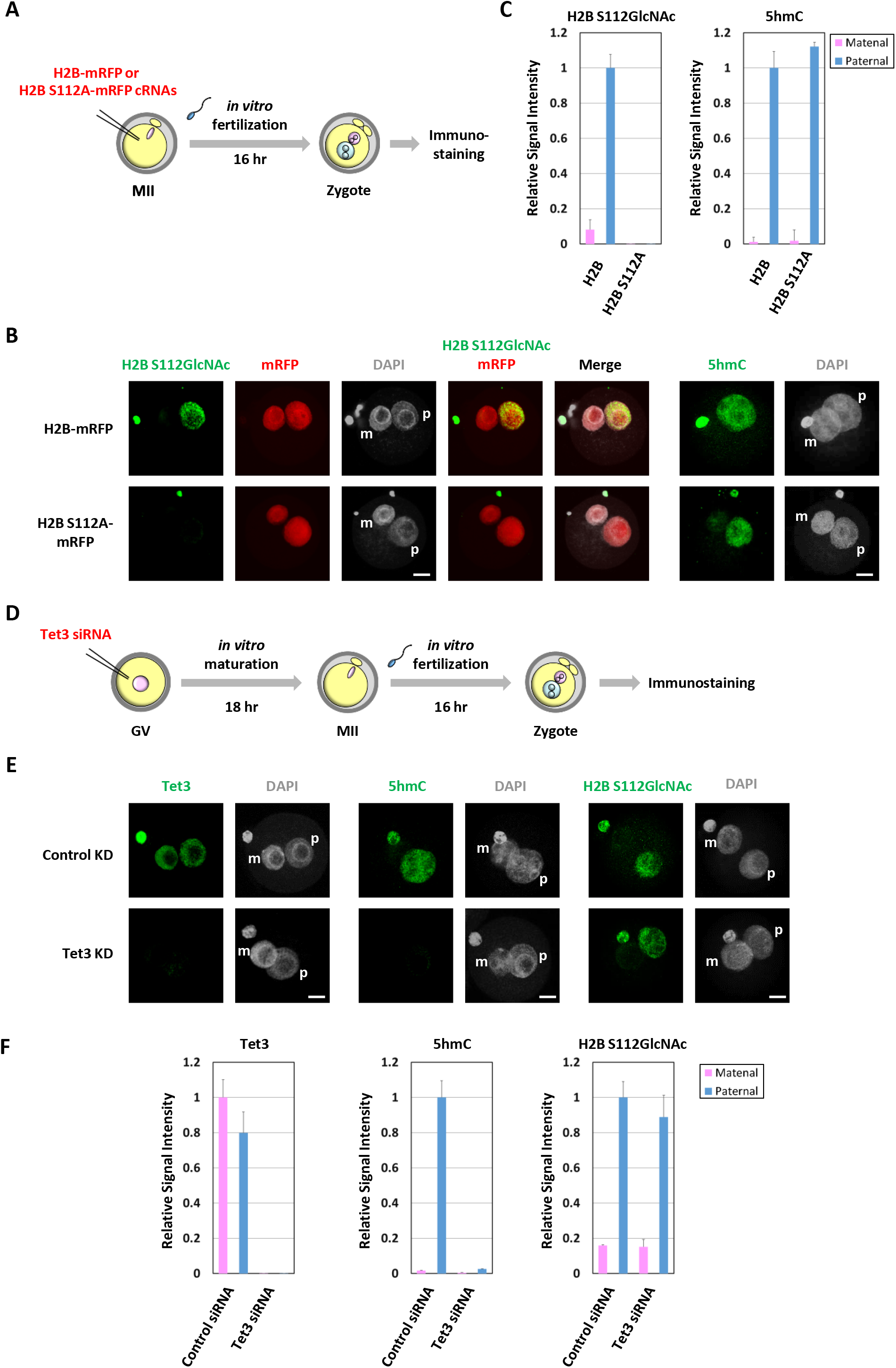
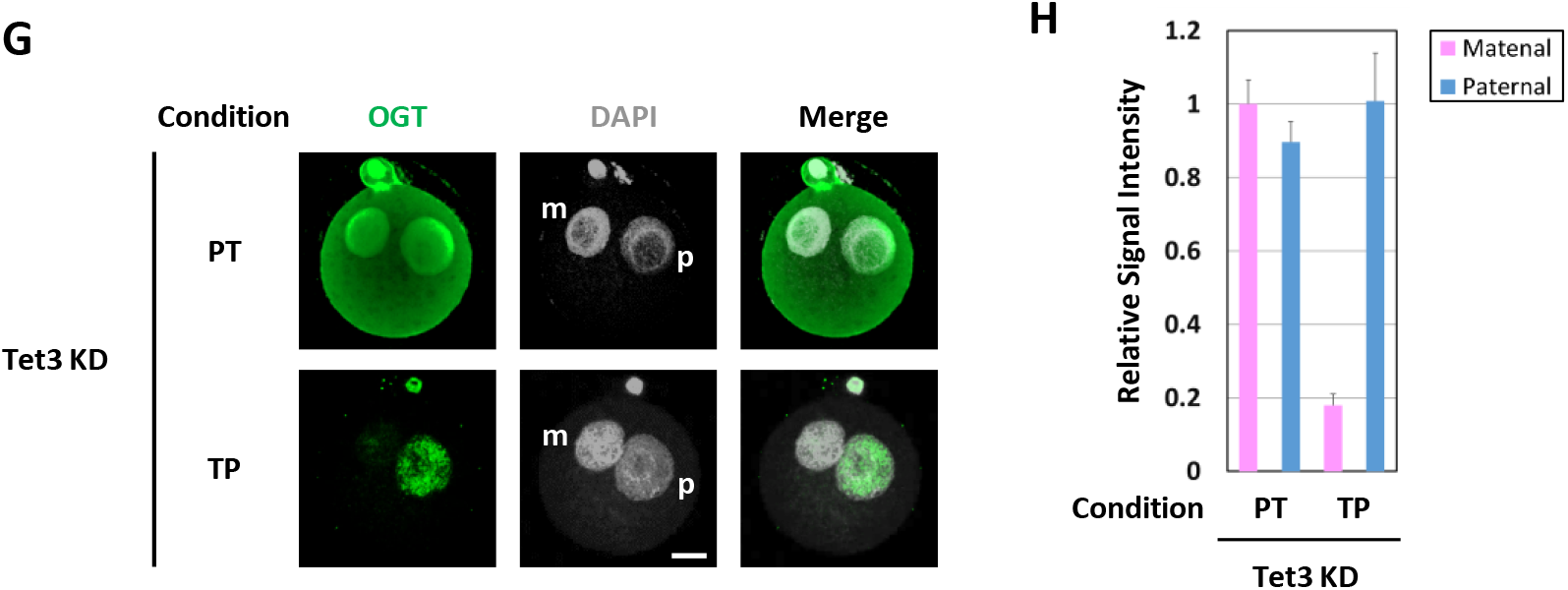
Tet3 and OGT were independently attached to paternal chromatin in zygote.t. (A) Schematic diagram of the preparation of H2B S112GlcNAc depleted zygotes. cRNAs of H2B-mRFP or H2B S112A-mRFP were microinjected into MII oocyte, and then fertilised *in vitro* and are cultured for 16 hr in potassium-enriched simplex optimized medium (KSOM). (B, C) H2B-mRFP or H2B S112A-mRFP expressing zygotes were stained with indicated antibodies (m, maternal pronuclei; p, paternal pronuclei). H2B S112GlcNAc and mRFP are shown in green and red, respectively (light panel); nuclei were stained with DAPI (grey). 5hmC is shown in green (right panel); nuclei were stained with DAPI (grey) (B). Twenty-two H2B-mRFP expressing zygotes and 28 H2B S112A-mRFP expressing zygotes were stained using anti-H2B S112GlcNAc antibody (light panel). Nineteen H2B-mRFP expressing zygotes and 25 H2B S112A-mRFP expressing zygotes were stained using anti-5hmC antibody (right panel). Intensities of H2B S112GlcNAc or 5hmC signals of parental chromatin were quantified using Image J (C). Error bars indicate standard deviation. d, Schematic diagram of the preparation of Tet3 knock down (KD) zygotes. siRNA specific for Tet3 (siTet3) or control siRNA (siControl) were microinjected into GV oocyte. After *in vitro* maturation, zygotes were obtained by *in vitro* fertilisation. (E, F) Control and Tet3 KD zygotes were stained with indicated antibodies. Tet3 (left panel), 5hmC (middle panel), and H2B S112GlcNAc (right panel) were shown in green; nuclei were stained with DAPI (grey) (E). Eleven control and 18 Tet3 KD zygotes were stained using anti-Tet3 antibody (light panel). Seventeen control and 19 Tet3 KD zygotes were stained using anti-5hmC antibody (middle panel). Twenty-three control and 18 Tet3 KD zygotes were stained using anti-H2B S112GlcNAc antibody (right panel). Intensities of Tet3 (light), 5hmC (middle), or H2B S112GlcNAc (right) signals of parental chromatin were quantified using Image J (F). Error bars indicate standard deviation. **g, h**, Tet3 KD zygotes were treated with PT or TP condition. Nineteen and 13 Tet3 KD zygotes were stained with OGT antibody under PT and TP conditions, respectively (m, maternal pronuclei; p, paternal pronuclei). OGT is shown in green; nuclei were stained with DAPI (grey) (G). Intensities of nuclear OGT signals of parental chromatin were quantified using Image J (G). Error bars indicate standard deviation. Scale bar, 20 μm.

We next generated Tet3 knockdown (KD) zygotes by injecting Tet3-targeting siRNA into GV-stage oocytes, followed by *in vitro* maturation and *in vitro* fertilization (Fig. 2D). As shown in Fig. 2E and F, endogenous Tet3 protein was completely depleted, and a significant reduction in paternal 5hmC was observed in Tet3 KD zygotes. Although Tet proteins have been reported to act as scaffolding factors that recruit OGT to chromatin, these Tet3 KD results indicate that Tet3 and OGT associate with paternal chromatin independently in zygotes. Supporting this notion, while OGT was predominantly detected in both maternal and paternal pronuclei under the PT fixation condition in Tet3 KD zygotes, OGT remained specifically localized to the paternal pronucleus under the TP condition (Fig. 2G, H).

### Concomitant chromatin modifications by OGT and Tet3 are required for minor zygotic genome activation

We next examined the effects of OGT and Tet3 on minor ZGA using a BrUTP incorporation assay (Fig. 3A). As shown in Fig. 3b and c, the transcriptional activity of the paternal genome was unchanged in both H2B S112GlcNAc-depleted zygotes and Tet3 KD zygotes. However, when H2B S112GlcNAc was depleted in Tet3 KD zygotes, paternal transcriptional activity was significantly reduced. These results indicate that concomitant chromatin modifications by OGT and Tet3 are prerequisite for minor ZGA in zygotes (Fig. 3B, C). Growing evidence suggests that Tet proteins are more versatile and multifaceted than initially appreciated, with important non-catalytic functions beyond cytosine hydroxymethylation (Krueger et al. 2017; Ketchum et al. 2024). We therefore performed a complementation assay to determine whether the enzymatic activity of Tet3—conversion of 5mC to 5hmC—is required for ZGA. Tet3-targeting siRNA designed against the 3’UTR was microinjected into GV-stage oocytes to deplete endogenous Tet3, and MII oocytes were obtained following in vitro maturation (Supplementary Fig. 4A). cRNAs encoding a Tet3 catalytic mutant together with mRFP-tagged H2B or its S112A mutant were subsequently microinjected into Tet3-depleted MII oocytes, and zygotes lacking H2B S112GlcNAc but expressing the Tet3 catalytic mutant were generated by *in vitro* fertilization (Supplementary Fig. 4A, B). The observed inhibition of ZGA in these zygotes demonstrated that the enzymatic activity of Tet3 is required for ZGA (Supplementary Fig. 5).

**Figure 3.**
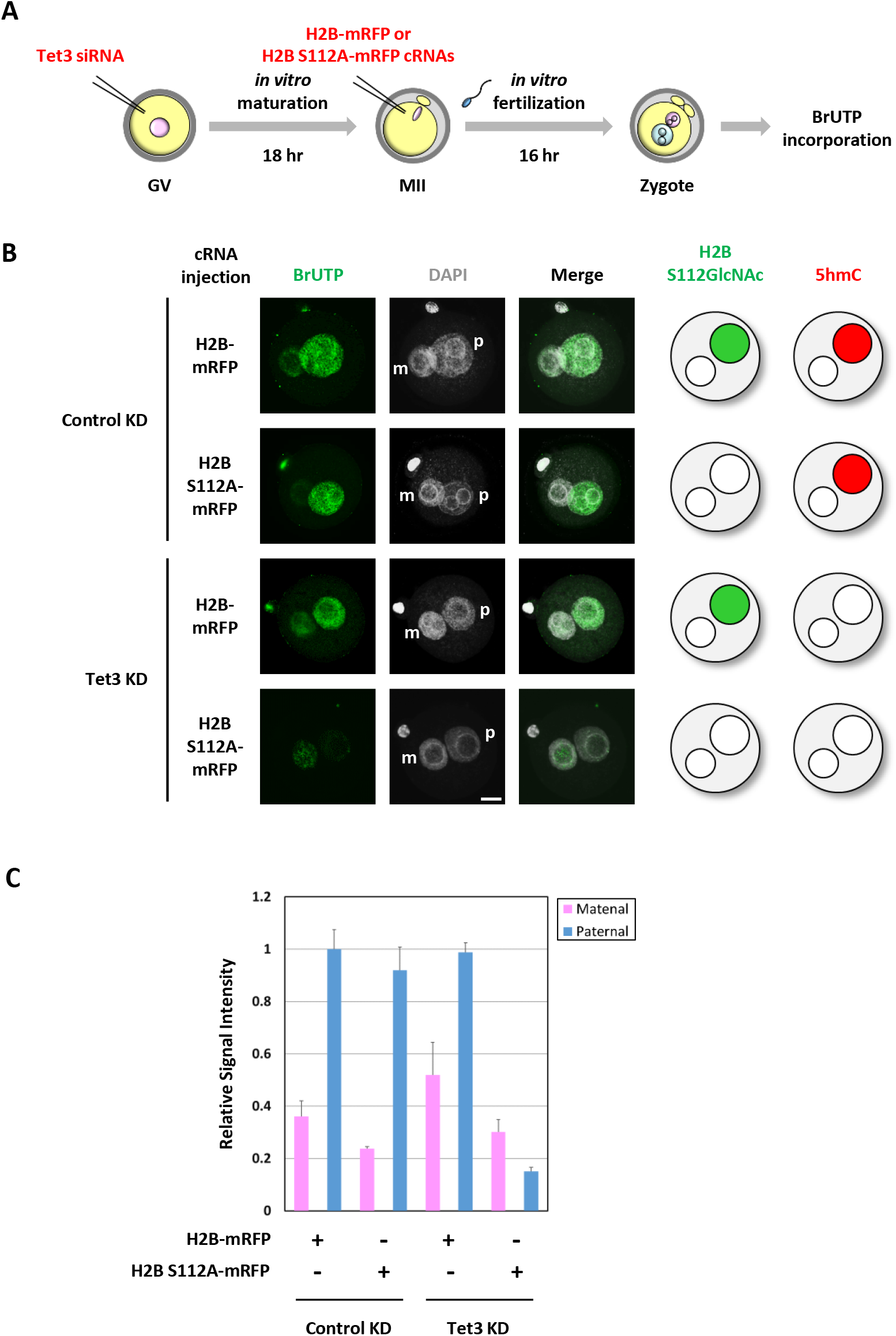
H2B S112GlcNAc and 5hmC in paternal chromatin are prerequisite for global transcriptional activation in zygotes.t. (A) Schematic diagram of the preparation of Tet3 and H2B S112GlcNAc depleted zygotes. Tet3 siRNA or scrambled control siRNA were microinjected into GV oocyte. After *in vitro* maturation, cRNAs of H2B-mRFP or H2B S112A-mRFP were microinjected, and Tet3 and H2B S112GlcNAc depleted zygotes were obtained by *in vitro* fertilisation. (B, C) BrUTP incorporation assay were performed using Tet3 and H2B S112GlcNAc depleted zygotes. Representative images of control, H2B S112-depleted, Tet3-depleted, and H2B S112GlcNAc and Tet3-depleted zygotes stained with anti-BrUTP antibody (m, maternal pronuclei; p, paternal pronuclei). BrUTP is shown in green; nuclei were stained with DAPI (grey) (B). Twenty-three H2B-mRFP expressing zygotes, 19 H2B S112A-mRFP expressing zygotes, 28 H2B-mRFP expressing Tet3 KD zygotes, and 38 H2B S112A-mRFP expressing Tet3 KD zygotes were stained using anti-BrUTP antibody. Intensities of BrUTP signals of parental pronuclei were quantified using Image J (C). Error bars indicate standard deviation. Scale bar, 20 μm.

### Zygotic genome transcription requires both Tet3 and H2B S112 O-GlcNAcylation

To examine the effects of 5hmC and H2B S112GlcNAc in more detail, we performed quantitative RT-PCR (qRT-PCR) analysis of zygotes depleted of Tet3 and/or H2B S112GlcNAc. Transcripts present in zygotes consist of maternally inherited RNAs as well as those newly transcribed from the zygotic genome. To specifically assess ZGA-derived transcripts, we selected the genes Dux, Gm8994, Zfp352, and Dub2a, all of which are transcribed in zygotes but not in MII oocytes, based on published datasets (Supplementary Fig. 6) (Park et al. 2015). Expression levels of these genes in Tet3 KD zygotes or in zygotes expressing the H2B S112A mutant alone were comparable to those in control zygotes (Fig. 4). In striking contrast, transcription of all analyzed genes was severely impaired in Tet3 KD zygotes expressing H2B S112A (Fig. 4). These results clearly demonstrate that both Tet3 and H2B S112GlcNAc are required for transcription from the zygotic genome (Supplementary Fig. 7).

**Figure 4.**
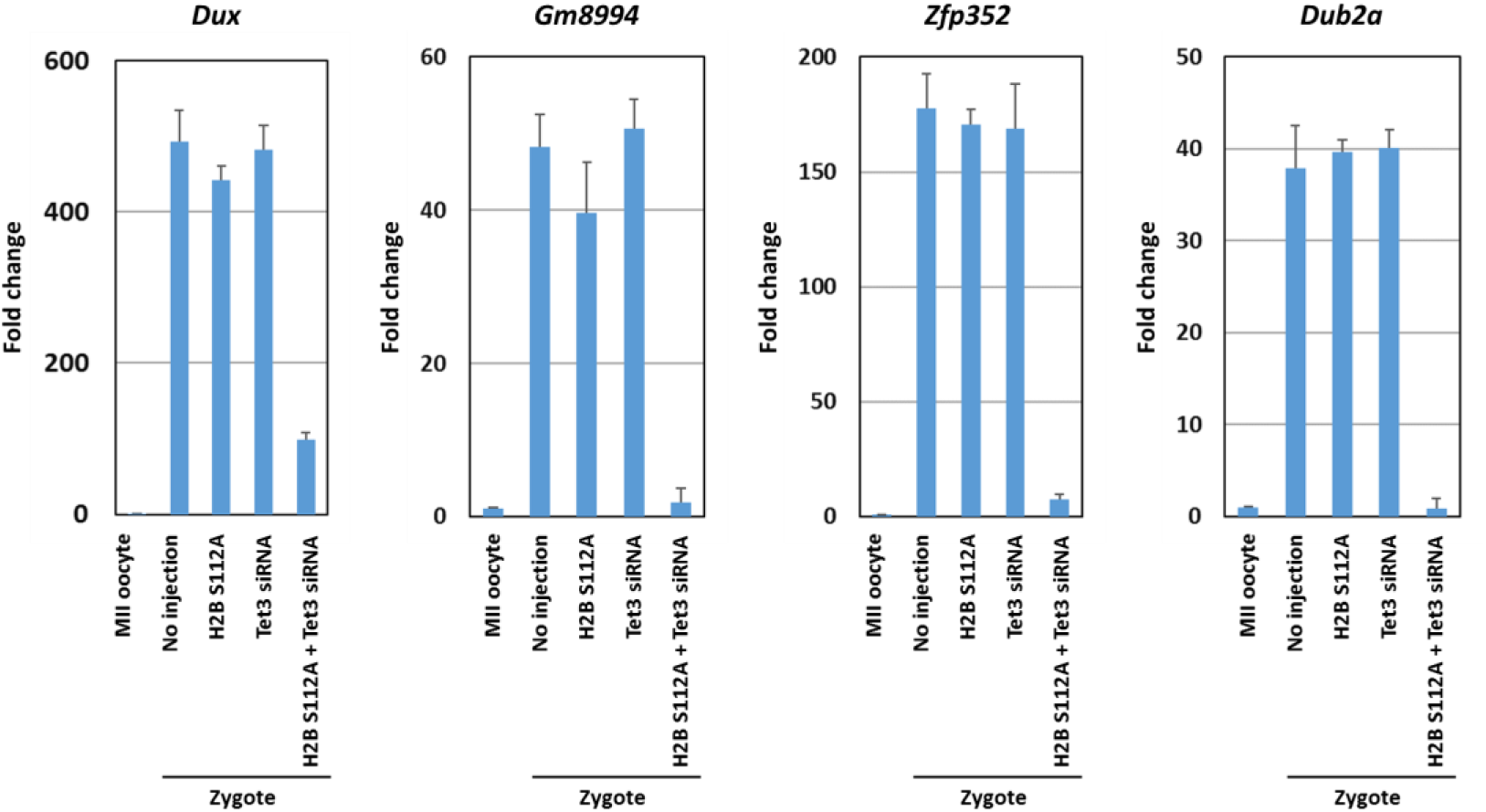
H2B S112GlcNAc and 5hmC are required for zygotic genome activatoin in zygotes.t. qRT-PCR analysis of Dux, Gm8994, Zfp352, and Dub2a were performed using cDNA synthesized from MII oocytes, control zygote, H2B S112A-mRFP expressing zygotes, Tet3 KD zygotes, and H2B S112A-mRFP expressing Tet3 KD zygotes. The relative expression level is normalized to internal control. Error bars indicate standard deviation.

Although minor ZGA is known to be critically important for the maternal–zygotic transition and subsequent embryonic development, the mechanisms underlying transcription in zygotes remain poorly understood. Nevertheless, several lines of circumstantial evidence suggest that chromatin structure plays a central role in regulating gene expression at this stage (Xu et al. 2025). Notably, actively expressed genes in zygotes lack canonical cis-regulatory elements in their upstream and downstream regions (Abe et al. 2015; Yamamoto et al. 2016). In addition, zygotes are thought to employ unique transcriptional regulatory mechanisms, as enhancers do not enhance transcriptional activity in this context (Majumder et al. 1993; Yamamoto and Aoki 2017). Furthermore, analyses of histone mobility in preimplantation embryos have revealed that zygotes possess the loosest chromatin structure among early developmental stages (Boskovic et al. 2014; Ooga et al. 2016). Given that OGT-mediated O-GlcNAcylation of histone H2B at Ser112 (H2B S112GlcNAc) facilitates H2B K120 monoubiquitination and promotes transcriptionally active chromatin in ES cells, it is reasonable to hypothesize that the same modification may contribute to establishing a permissive chromatin environment during ZGA. Because ZGA requires a rapid shift toward an open chromatin state and robust transcriptional activation of the embryonic genome, H2B S112GlcNAc is likely to support this process by promoting chromatin accessibility and facilitating transcription-related histone crosstalk, particularly through H2B K120 ubiquitylation. The nutrient-responsive nature of O-GlcNAcylation further suggests that H2B S112GlcNAc could act as a metabolic integrator that helps coordinate the onset of ZGA with maternal metabolic cues. Although chromatin accessibility was not directly measured in this study, our findings, together with previous reports of unusually high histone mobility in zygotes, support the model that coordinated DNA and histone modifications facilitate chromatin relaxation during minor ZGA.

## Materials and methods

### Plasmids

The H2B S121A-mRFP was generated PCR-based mutagenesis and confirmed by sequencing. The primer used to generate the mutant are described in Supplementary Table 1.

### Zygote collection and culture

Female BDF1, or Stella KO mice >8 weeks old were superovulated by injecting 7.5 U of human chorionic gonadotropin (hCG; ASKA pharmaceutical) 48 h after injecting 7.5 U of pregnant mare serum gonadotropin (PMSG; ASKA pharmaceutical), and then mated with male BDF1 mice. Fertilised eggs were collected from the oviduct, placed in 100-μL drops of potassium simplex optimization medium (KSOM; Millipore), and cultured at 37°C in an atmosphere of 5% CO_2_. All experiments were carried out according to the guidelines of the Committee on the Ethics of Animal Experiments of the Nagahama Institute of Bio-Science and Technology and the Osaka University Animal Care and Use Committee.

### Generation of Tet3 KD zygote

Fully grown germinal vesicle (GV) oocytes were obtained from 4 - 8 weeks old female BDF1 mice 48 h after injection with 7.5 U of PMSG. The ovaries were removed from the mice ant transferred to M2 media (Sigma-Aldrich) containing 0.2 mM 3-isobutyl-1-methylxanthine (IBMX; Sigma-Aldrich). The ovarian follicles were punctured with a 27-guage needle, and the cumulus cells were removed from cumulus-oocyte complexes using a narrow-bore glass pipette. The GV-oocytes were transferred into α-MEM (Wako) supplemented with 5% fatal calf serum (FCS; Sigma), and 10 ng/mL epidermal growth factor (EGF; Wako). After 1h of incubation, GV-oocytes were microinjected with 3-5 pL of 2M scrambled control siRNA, Tet3 siRNA, or Tet3 3’UTR siRNA with FemtoJet 4i (Eppendrf). After 1h of microinjection, GV-oocytes were washed with IBMX-free α-MEM supplemented with 5% FCS and 10 ng/mL EGF, and cultured for 18 h at 37°C in an atmosphere of 5% CO_2_ to obtain MII oocytes. And then subject to *in vitro* fertilisation (IVF) to obtain control, Tet3 KD-zygote. siRNA sequences used in this study are listed in Supplementary Table 1.

### In vitro transcription

Capped mRNA was synthesized using a T7 mMessage mMachine kit (Thermo Fisher Scientific). Poly (A) tails were added to the capped mRNA using a Poly (A) Tailing Kit (Thermo Fisher Scientific) according to the manufacturer’s protocol. The synthesized mRNAs were subjected to gel filtration using NucAway Spin Columns (Invitrogen) to remove unreacted substrates and then stored at -80°C until use.

### Generation of H2B-mRFP and H2B S112A-mRFP expressing zygote

Female BDF1 mice 4 - 8 weeks old were superovulated by injecting 7.5 U of hCG 48 h after injecting 7.5 U of PMSG, and then obtained Metaphase II (MII) oocytes 14 h after injecting hCG. The cumulus cells were removed by a short incubation in M2 medium (Sigma-Aldrich) containing 35 U/mL hyaluronidase (Sigma-aldrich), and washed with M2 medium, and microinjected 3-5 pL of 75 ng /mL H2B-mRFP or H2B S112A-mRFP cRNA with siRNA with FemtoJet 4i. And then subject to IVF to obtain H2B-mRFP or H2B S112A-mRFP expressing zygotes.

### in vitro fertilisation (IVF)

MII oocytes were transferred to Human Tubal Fluid (HTF; Millipore), and inseminated with capacitated spermatozoa. Spermatozoa were collected from the cauda epididymis of adult B6D2F1 male mice and incubated in HTF for 1 h. After 2h of insemination, obtained zygotes were transferred to KSOM and cultured for 16 h at 37°C in an atmosphere of 5% CO_2_.

### Triton treatment of zygotes

Zygotes were treated with Triton X-100 to a previous report. Zygotes were treated with 0.2% Triton X-100 in PBS for 45-60 s until the perivitelline space was eliminated. Immediately after Triton treatment, the zygotes were washed with PBS at least five times and then fixed in 4% PFA. After washing with PBS, immunostaining was performed as described below.

### Immunohistochemistry

Fertilised eggs were washed with PBS, fixed for 15 min in 4% PFA in PBS at room temperature, and permeabilised with 0.2% Triton X-100 in PBS for 20 min at room temperature. The eggs were blocked for 1 h in 5% normal goat serum (NGS; Sigma-aldrich) in PBS at room temperature and incubated overnight at 4°C with primary antibodies as shown in Supplementary Table 2. The following day, the eggs were washed three times with 0.05% Tween20 in PBS, and staining was detected by incubating the eggs with secondary antibodies as shown in Supplementary Table 1. Nuclei were stained with 1 μg/mL DAPI. 5hmC staining was performed as described previously (Nakamura et al. 2012). Immunofluorescence was visualised using an FV10i confocal laser scanning microscope (Olympus).

### BrUTP incorporation assay

BrUTP incorporation assay was performed as described previously (Aoki et al. 1997). Zygotes were washed with PBS, and treated with 0.05 % Triton X in physiological buffer (PB) containing 100 mM potassiu m acetate, 30 mM KCl, 1 mM MgCl_2_, 10 mM Na_2_HPO_4_, 1 mM ATP supplemented with 1 mM DTT, 0.2 mM phenylmethylsulfonyl fluoride, and 50 U/mL RNasin (Promega) for 1 - 2min to permeabilize the plasma membrane. Zygotes were washed with PB, and then transferred to 100 mM potassium acetate, 30 mM KCl, 1 mM MgCl_2_, 10 mM Na_2_HPO_4_ containing 2 mM ATP, 0.4 mM each of GTP, CTP, and BrUTP, and 2 mM MgCl_2_, and incubated for 15 min at 33°C. Zygotes were washed with PB and then treated with PB containing 0.2% Triton X-100 for 3min at room temperature. After incubation, zygotes were washed with PB, and then fixed with 4% PFA/PBS for 15 min, and subject to immunohistochemistry.

### Quantative reverse transcription real-time PCR analysis

Total RNA was purified with RNeasy Micro Kit (Qiagen) according to the manufacturer’s protocol. As an external control, 10 pg of λ polyA^+^ RNA (Takara) was added prior to total RNA purification. cDNA was synthesized using SuperScript VILO cDNA Synthesis Kit (Invitrogen) according to the manufacturer’s protocol. Real-time PCR were performed on LigtCycler 480 Instrument II (Roche) using KAPA SYBR FAST qPCR Master Mix (2X) Kit (KAPA Biosystems). The relative expression levels of each gene were normalized to those of external control. The primers used in this study are listed in Supplementary Table 1.

### Co-immunoprecipitation assay

Co-immunoprecipitation assay was performed as described previously (Nakamura et al. 2007).

## Competing interest statement

All authors declare no competing interests.

## Acknowledgements

We thank Drs. K.Yamagata and R.Fujiki for providing H2B-mRFP expression plasmid and anti-histone H2B S112GlcNAc antibody, respectively. This work was supported in part by a Gran-i-Aid for Scientific Research on Research (B) (#25291053) to T. Nakamura from MEXT/JSPS, by a Grant-in-Aid for Scientific Research on Innovative Areas (‘Analyses and regulation of germline epigenome’) (#25112006) to T. Nakamura from MEXT/JSPS, and by Core Research for Evolutional Science and Technology (CREST) (#J150701424) from AMED, Japan, to T.Nakamura and T. Nakano.

## Author contributions

T.Nakamura and T.Nakano conceived the project and wrote the manuscript. T.Nakamura designed, performed the all experiments and evaluated the results. A.F. performed some experiments with T.Nakamura. T.Nakatani propagated Stella KO mice and provided zygote from Stella KO female mice.

## Author Information

Correspondence and requests for materials should be addressed to T. Nakamura

## Notes

### Competing Interest Statement

The authors have declared no competing interest.

