## Supplementary figures for "Concomitant DNA hydroxymethylation and histone H2B O-GlcNAcylation are prerequisites for zygotic genome activation in mice"

Nakamura T et al._Supplementary Figure 1

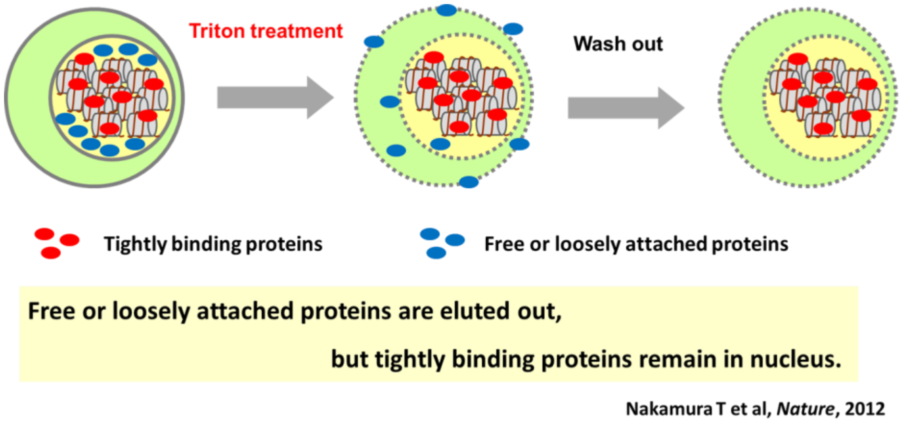

Nakamura T et al._Supplementary Figure 2

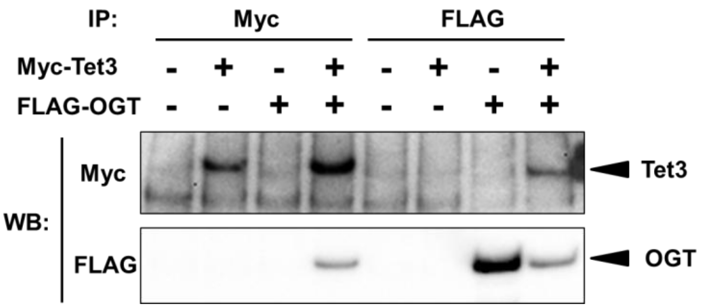

Nakamura T et al._Supplementary Figure 3

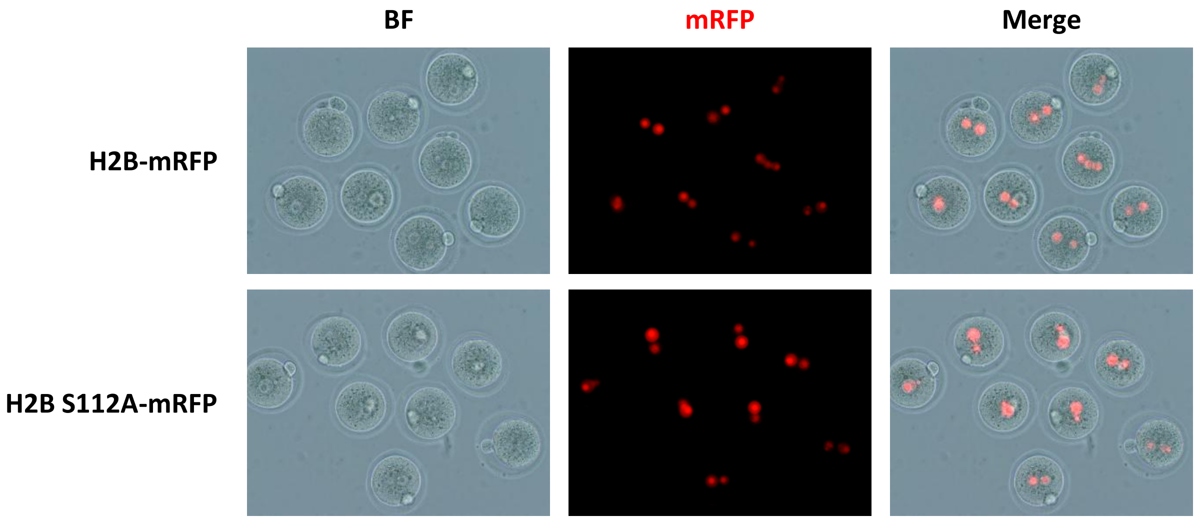

Nakamura T et al._ Supplementary Figure 4

A

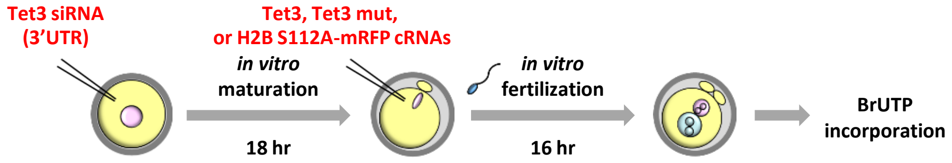


B

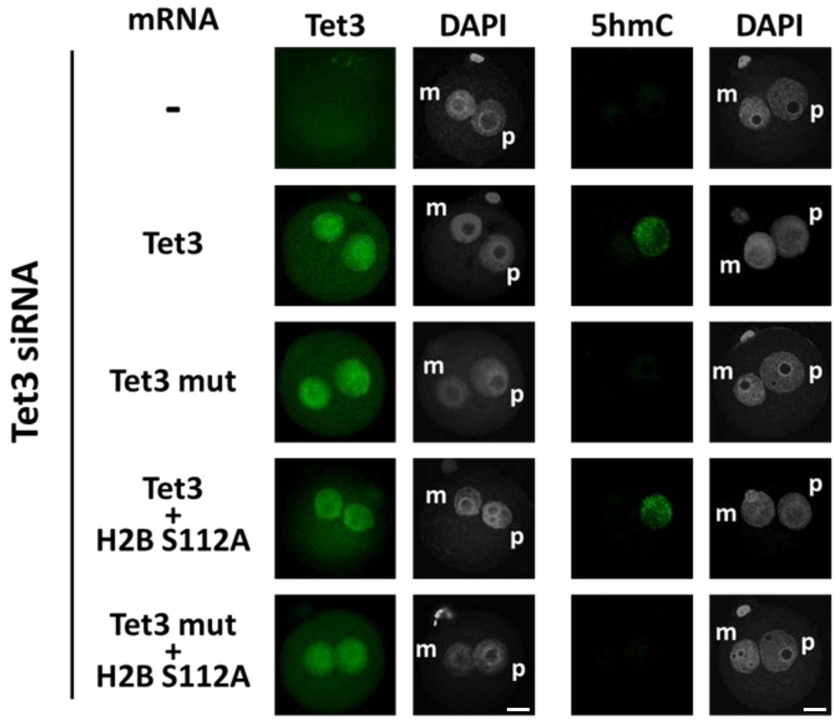

Nakamura T et al._ Supplementary Figure 5

**A**

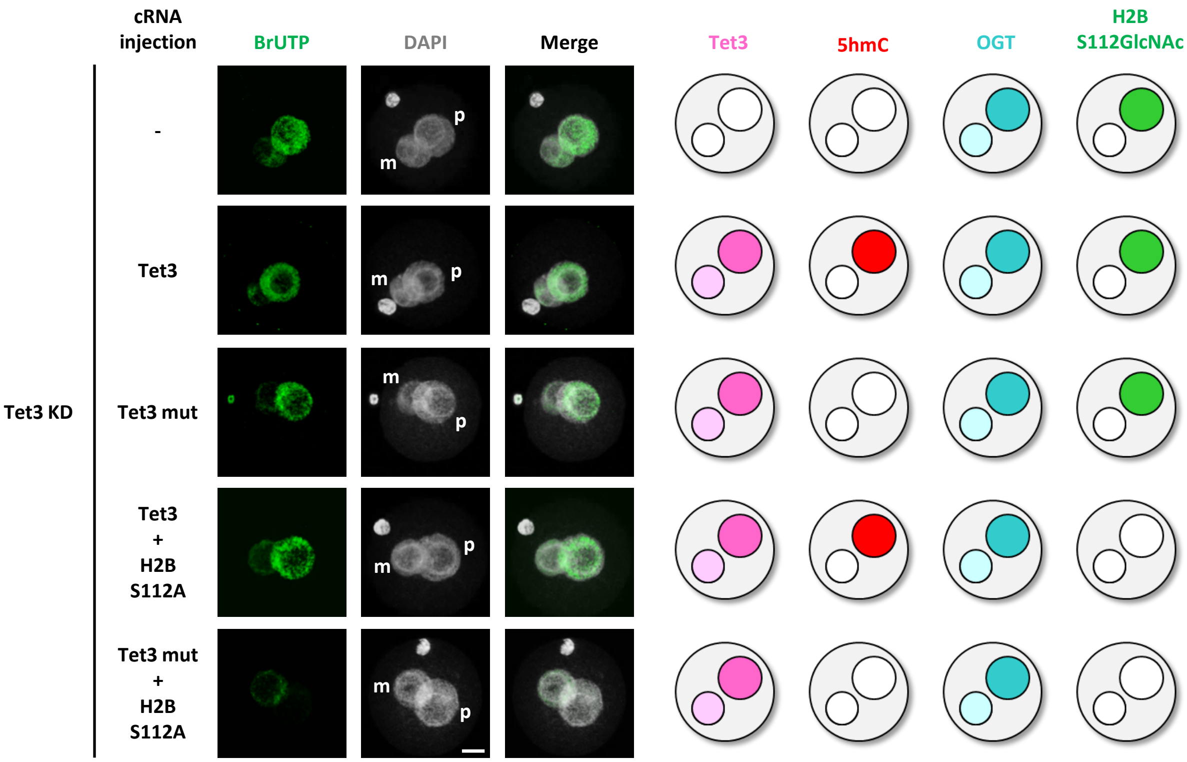


**B**

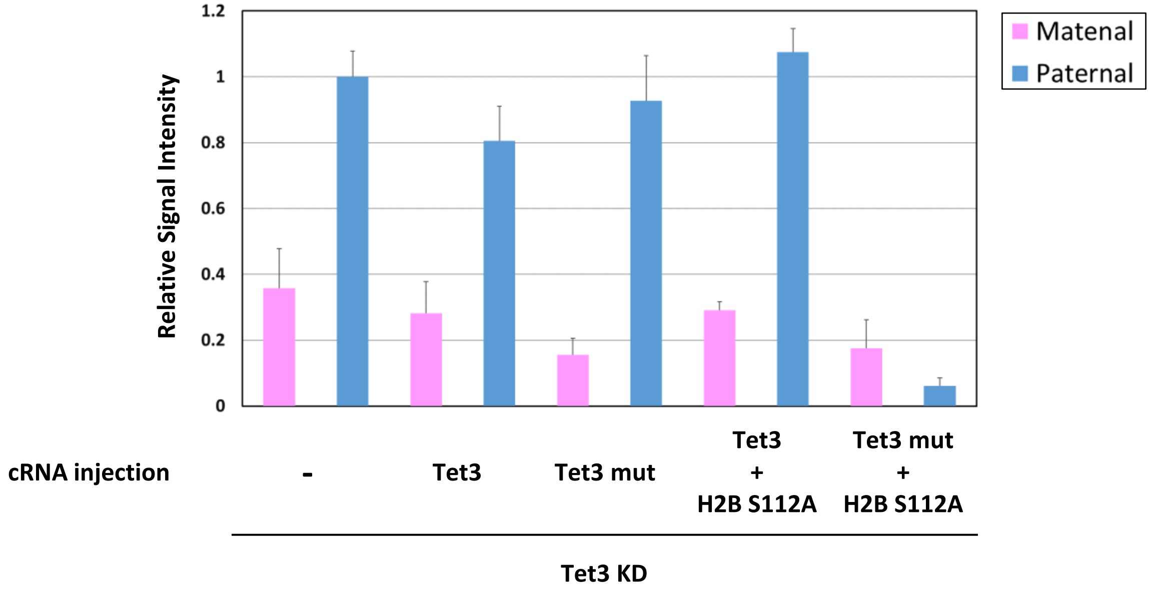

Nakamura T et al._ Supplementary Figure 6

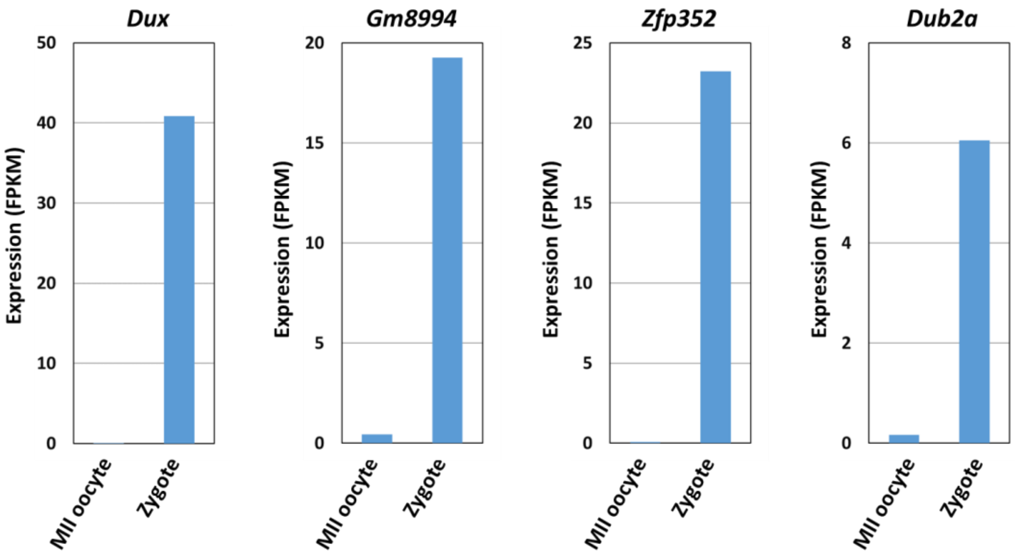

Nakamura T et al._ Supplementary Figure 7

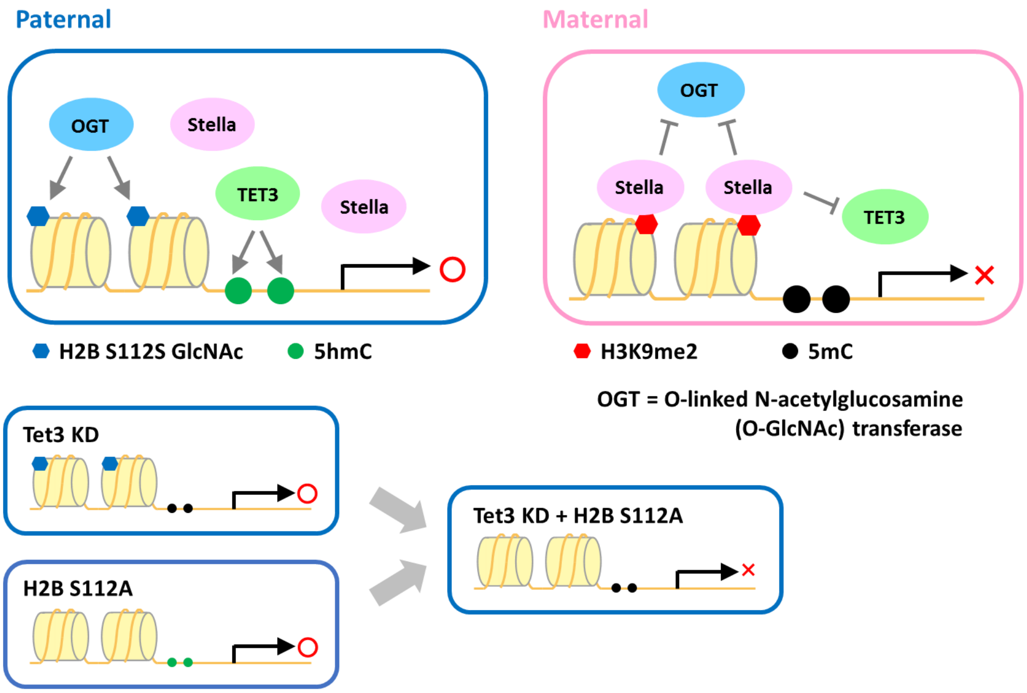
