## Supplementary figure legends for "Concomitant DNA hydroxymethylation and histone H2B O-GlcNAcylation are prerequisites for zygotic genome activation in mice"

**Supplementary figure 1. Evaluation of the binding affinity of nuclear proteins to chromatin.**
Pre-extraction with Triton X-100 prior to PFA fixation removes nuclear proteins that are unbound or weakly associated with chromatin.

**Supplementary figure 2. Binding of Tet3 to OGT.**
Myc-tagged Tet3 was coexpressed with FLAG-tagged OGT in 293T cells. Tet3 and OGT were immunoprecipitated with anti-FLAG and anti-Myc antibodies, respectively. The immunoprecipitates were analyzed by western blotting.

**Supplementary figure 3. Exogenous histone H2B and its mutant are efficiently incorporated into both pronuclei of the fertilized egg.**
cRNA encoding H2B-mRFP and H2B S112A-mRFP was microinjected into MII oocytes, followed by in vitro fertilization (IVF).

**Supplementary figure 4. Effect of Tet3 enzymatic activity on the conversion of 5mC to 5hmC.**
GV-stage oocytes were microinjected with Tet3 3' UTR siRNA and subjected to IVM. The resulting MII oocytes were further injected with the indicated cRNAs (Tet3, Tet3 mut, and/or H2B S112A), followed by IVF to obtain zygotes. The status of 5hmC was then analyzed at the pronuclear stage.

**Supplementary figure 5. Effect of Tet3 enzymatic activity on BrUTP incorporation in zygotes.**
GV-stage oocytes were microinjected with Tet3 3' UTR siRNA and subjected to IVM. The resulting MII oocytes were further injected with the indicated cRNAs (Tet3, Tet3 mut, and/or H2B S112A), followed by IVF to obtain zygotes. BrUTP incorporation was then analyzed at the pronuclear stage to assess transcriptional activity.

**Supplementary figure 6. Expression profiles of minor ZGA-associated genes in MII oocytes and zygotes.**
Minor ZGA-associated genes were identified using the DBTMEE database.

**Supplementary figure 7. Schematic model for the regulation of minor ZGA by DNA hydroxymethylation and histone H2B O-GlcNAcylation.**

(Top) Asymmetric epigenetic regulation in paternal and maternal pronuclei. In the paternal pronucleus (blue box), TET3-mediated DNA hydroxymethylation (5mC to 5hmC) and OGT-mediated histone H2B S112 O-GlcNAcylation promote minor ZGA. In contrast, in the maternal pronucleus (pink box), Stella prevents these modifications and maintains H3K9me2 and 5mC, leading to transcriptional repression. (Bottom) Synergistic requirement of Tet3 and H2B O-GlcNAcylation for minor ZGA. Loss of either Tet3 (Tet3 KD) or H2B O-GlcNAcylation (H2B S112A) alone allows minor ZGA to proceed (indicated by red circles). However, the simultaneous loss of both Tet3 and H2B O-GlcNAcylation (Tet3 KD + H2B S112A) results in the failure of minor ZGA (indicated by the red X), demonstrating their redundant or synergistic role in activating the embryonic genome.
