## Supplementary table 1 for "Concomitant DNA hydroxymethylation and histone H2B O-GlcNAcylation are prerequisites for zygotic genome activation in mice"

Nakamura T et al. Supplementary Table S1 Primer sequences

| Name | Sequence (5' to 3') | Application |
|---|---|---|
| Tet3_catalytic mutant_FWD | AGGCCCAACATAACCTCTACAAT | mutagenesis |
| Tet3_catalytic mutant_REV | TGTGGGCGTGGGCACAGAAGTCC | mutagenesis |
| H2BS112S_FWD | GCCAAGCACGCCGTGGCCGAGGGTACTAAG | mutagenesis |
| H2BS112S_REV | CTTAGTACCCTCGGCCACGGCGTGCTTGGC | mutagenesis |
| Dux_FWD | CCCAGCGACTCAAACTCCTTC | RT-qPCR |
| Dux_REV | GGACTTCGTCCAGCAGTTGAT | RT-qPCR |
| Gm8994_FWD | CTCCAACCAGAGAGTTAGCGG | RT-qPCR |
| Gm8994_REV | CCCGTGTCGTTAAACTTCTCC | RT-qPCR |
| Zfp352_FWD | AAGTCCCACATCTGAAGAAACAC | RT-qPCR |
| Zfp352_REV | GGGTATGAGGATTCACCCACA | RT-qPCR |
| Dub2a_FWD | GGTAGTGGTGGAGCTAACTGC | RT-qPCR |
| Dub2a_REV | GACTCTGGGTTACATGGGCTT | RT-qPCR |
