## Supplementary table 2 for "Concomitant DNA hydroxymethylation and histone H2B O-GlcNAcylation are prerequisites for zygotic genome activation in mice"

Nakamura T et al. Supplementary Table S2 Antibodies

| Name | Dilution | Applicatoin |
|---|---|---|
| Anti-TET3 antibody (abcam; ab153724) | 1:200 | IF |
| Anti-H3K9me2(upstate; 07-441) | 1:200 | IF |
| Anti-H3K9me2 (abcam; ab1220) | 1:200 | IF |
| Anti-5hmC (Active motif; 39770) | 1:2000 | IF |
| Anti-BrdU (Merk; B8434) | 1:500 | IF |
| Anti-H2B S112GlcNAc (gifted from R. Fujiki) | 1:200 | IF |
| Anti-O-GlcNAc transferase (Santa Cruz; sc-74547) | 1:100 | IF |
| Anti-rabbit IgG-Alexa488 (Invitrogen; 11034) | 1:200 | IF |
| Anti-mouse IgG-Alexa488 (Invitrogen; 11029) | 1:200 | IF |
| Anti-rabbit IgG-Alexa568 (Invitrogen; 11036) | 1:200 | IF |
| Anti-mouse IgG-Alexa568 (Invitrogen; 11031) | 1:200 | IF |
| Anti-FLAG (Sigma; F3165) | 1:1000 | WB |
| Anti-Myc (Santa Cruz; sc-40) | 1:200 | WB |
| Anti-FLAG (Sigma; F3165) | 1 μg | IP |
| Anti-Myc (Santa Cruz; sc-40) | 1 μg | IP |

Abbreviation: IF, immuno fluorescence; WB, western blotting; IP, immunoprecipitation.
